# Environmental Consistency, Not Certainty, Governs Predictive Motor Strategies During Walking

**DOI:** 10.1101/2025.09.03.674073

**Authors:** Francis M. Grover, Anna Shafer, Keith E. Gordon

## Abstract

Humans rely on predictive control to maintain stability while walking in dynamic environments, yet the strategies driving this control remain unclear when environments are inconsistent or uncertain. We tested how people adapt to repeated lateral force field disruptions of switching directions, manipulating the force field’s Consistency (how often switches occurred) and Certainty (how predictable the switches were). We quantified motor adaptation (error reduction) and sought evidence for two predictive strategies: pattern prediction (detecting a global trial pattern) and carryover prediction (assuming the next trial matches the previous).

Adaptation profiles revealed a surprising finding: Consistency, not certainty, drove predictive control. When trials were Consistent, participants pursued appropriate predictive strategies (pattern or carryover). However, when trials were Inconsistent, participants exhibited counterproductive strategies, such as carryover prediction (despite the next trial most often being opposite) or no prediction (despite full certainty of each forthcoming trial). Error reduction was likewise dominated by consistency, with certainty exerting negligible influence.

These findings, counter to long-held observations in upper-limb control, suggest the central nervous system faces unique challenges in whole-body control of walking and cannot rely on the simpler adaptation strategies observed in isolated limb control. This raises new, important questions for control of walking and whole-body motor adaptation.

## 1. Introduction

### 1.1 Background

Maintaining walking balance in changing environments often depends on predictive control strategies that use prior sensorimotor experience to guide future movements. In consistent environments, people adopt predictive movements that compensate for expected environmental properties and reduce motor error (Bucklin et al., 2019; Shadmehr & Mussa-Ivaldi, 1994). In frequently changing environments, however, predictive strategies are only useful when changes are sufficiently cued or predictable (Howard et al., 2013; Hirashima & Nozaki, 2012). When changes are unpredictable, prediction is both unfeasible and costly, as incorrect predictions increase motor error (Shadmehr et al., 2010).

In such cases, people might abandon prediction altogether, either due to hesitation (i.e., choosing to overlook errors rather than try to learn from them; Albert et al., 2021; Fernandes, Stevenson, & Kording, 2012; Havermann & Lappe, 2010; Herzfeld et al., 2014) or instead to pursue protective strategies (e.g., impedance or averaging; Burdet et al., 2001; Darainy & Ostry, 2008; Gribble et al., 2003; Scheidt et al., 2001) that enhance stability in all directions and, while not as efficient as a successful predictive strategy (Hogan, 1985), still enables some decreases in motor error (Franklin et al., 2003). However, prior work shows that even in unpredictable walking environments, people still attempt predictive strategies. Specifically, they assume upcoming disruptions will resemble previous ones, a counterproductive bias we term **carryover prediction** (Bucklin et al., 2023).

This raises new questions about predictive control even in consistent environments. Consistent predictive responses to a consistent force field (Bucklin et al., 2019) could reflect either the same carryover prediction seen under randomness (Bucklin et al., 2023), or alternatively that participants identified a global pattern—that all applied forces were in the same direction (what we label **pattern prediction**). In a consistent environment, both strategies would produce the same behavior.

These recent studies therefore motivate a need to more precisely understand when and which (carryover or pattern) predictive strategies are pursued by individuals during walking, and how this affects overall sensorimotor adaptation (error reduction). This distinction becomes especially important when environments vary along two dimensions: *certainty* (ability to predict the next trial) and *consistency* (how often conditions change across trials). For example, a fixed alternating trial sequence (e.g., reversing force fields) is low in consistency but high in certainty, supporting predictive control only if people engage in pattern prediction and not carryover prediction.

To disentangle these strategies, certainty and consistency must be manipulated independently, enabling direct of: **1) Under which conditions predictive control is pursued, and which form it takes (carryover or pattern)**, and **2) How the varying pursuit of each strategy affects overall adaptation and error reduction.**

### 1.2 Current Study

To address these two goals, healthy adults performed repeated stepping trials while exposed to a lateral **Force Field** directed either rightward or leftward to their pelvis. We tested different trial schedules that varied in Force Field direction consistency and certainty. **Consistent** schedules contained long stretches where Force Field direction persisted across trials (**Streaks**), whereas **Inconsistent** schedules exhibited a high frequency of **Switches** in Force Field direction. Within each, **Certain** schedules followed a fixed pattern that participants were informed of, while **Uncertain** schedules included small probabilistic deviations of which participants were not informed. This yielded four experimental groups: **Consistent-Certain**, **Inconsistent-Certain**, **Consistent-Uncertain**, and **Inconsistent-Uncertain** (Fig. 1).

**Figure 1.**
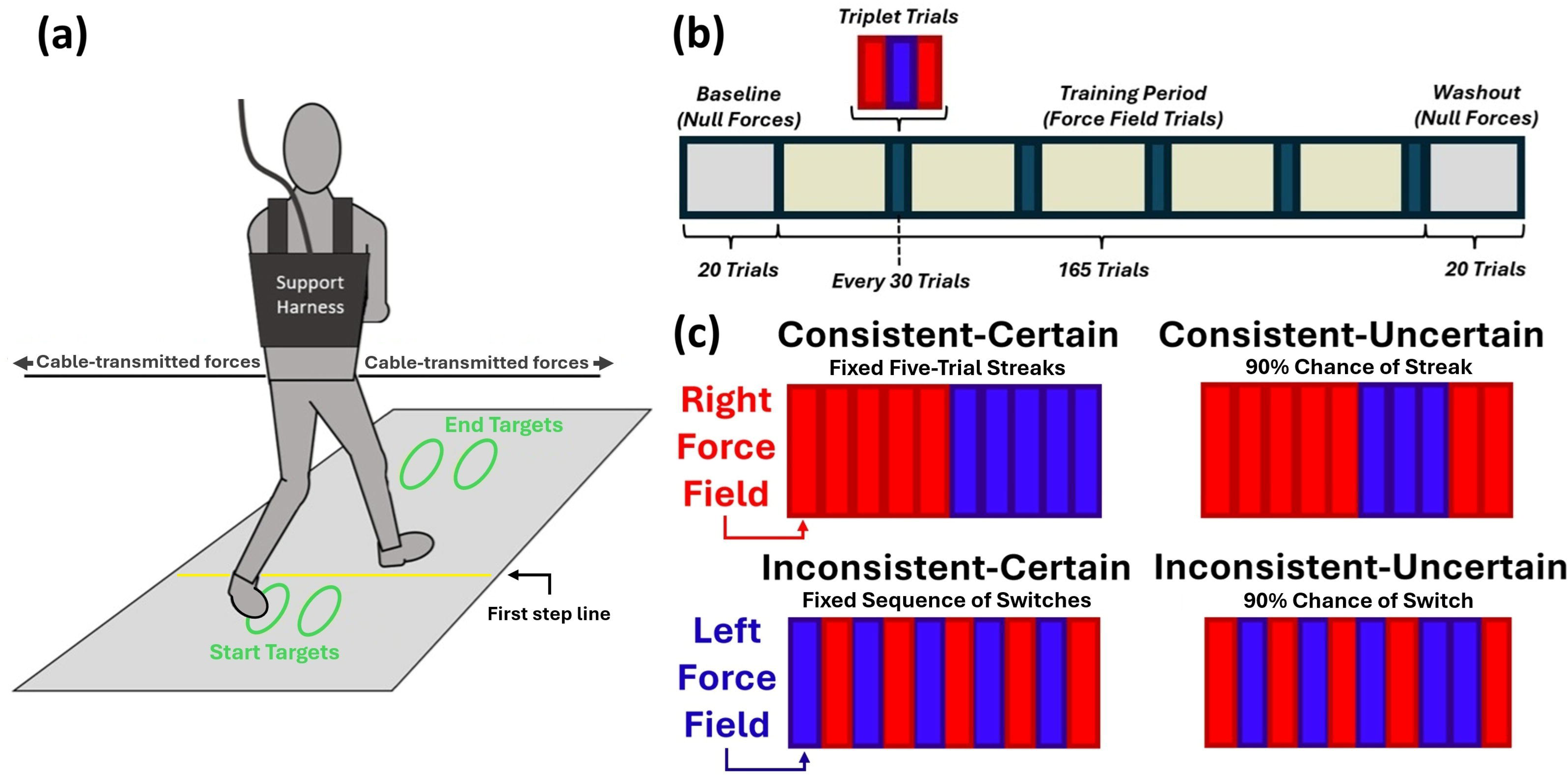
(a) Experimental set-up and discrete stepping paradigm. On each trial, participants performed three steps from the start to end targets while a cable-driven device applied a lateral Force Field (rightward or leftward) to the pelvis. (b) Protocol. All participants completed Baseline (Null Forces), Training Period with scheduled triplet sequences (Right-Left-Right), and Washout (Null Forces). (c) Trial schedules. Consistent groups experienced streaks of the same direction; Inconsistent groups experienced frequent switches. Certain groups followed fixed, informed schedules, whereas Uncertain groups followed probabilistic, uninformed schedules.

We first quantified overall error reduction across training, then examined how motor errors fluctuated on Switch versus Streak trials specifically to gain insight into the control strategies participants employed. To more directly investigate these specific motor strategies, we also assessed **Triplet Trials,** three trial sequences during which the Force Field direction follow a fixed progression (Right, Left, Right), to quantify how directional bias in early movements (i.e., bias in movements prior to Force Field onset) changed across each direction Switch.

### 1.3 Hypotheses

Previous studies (Bucklin et al., 2019; 2023; Jiang & Grover et al. 2024) showed that, when adapting to a consistent Force Field, participants bias early movement in a specific direction to pre-emptively resist it. We therefore based our hypotheses about predictive strategies around this bias in early movement. Pattern prediction would be indicated by biases aligned with the overall sequence of Force Field directions, whereas carryover prediction would be indicated by biases aligned with the immediately preceding trial. For details on how these hypotheses were operationalized, see Section 2.6 (Analyses) and Figure 2.

**Figure 2.**
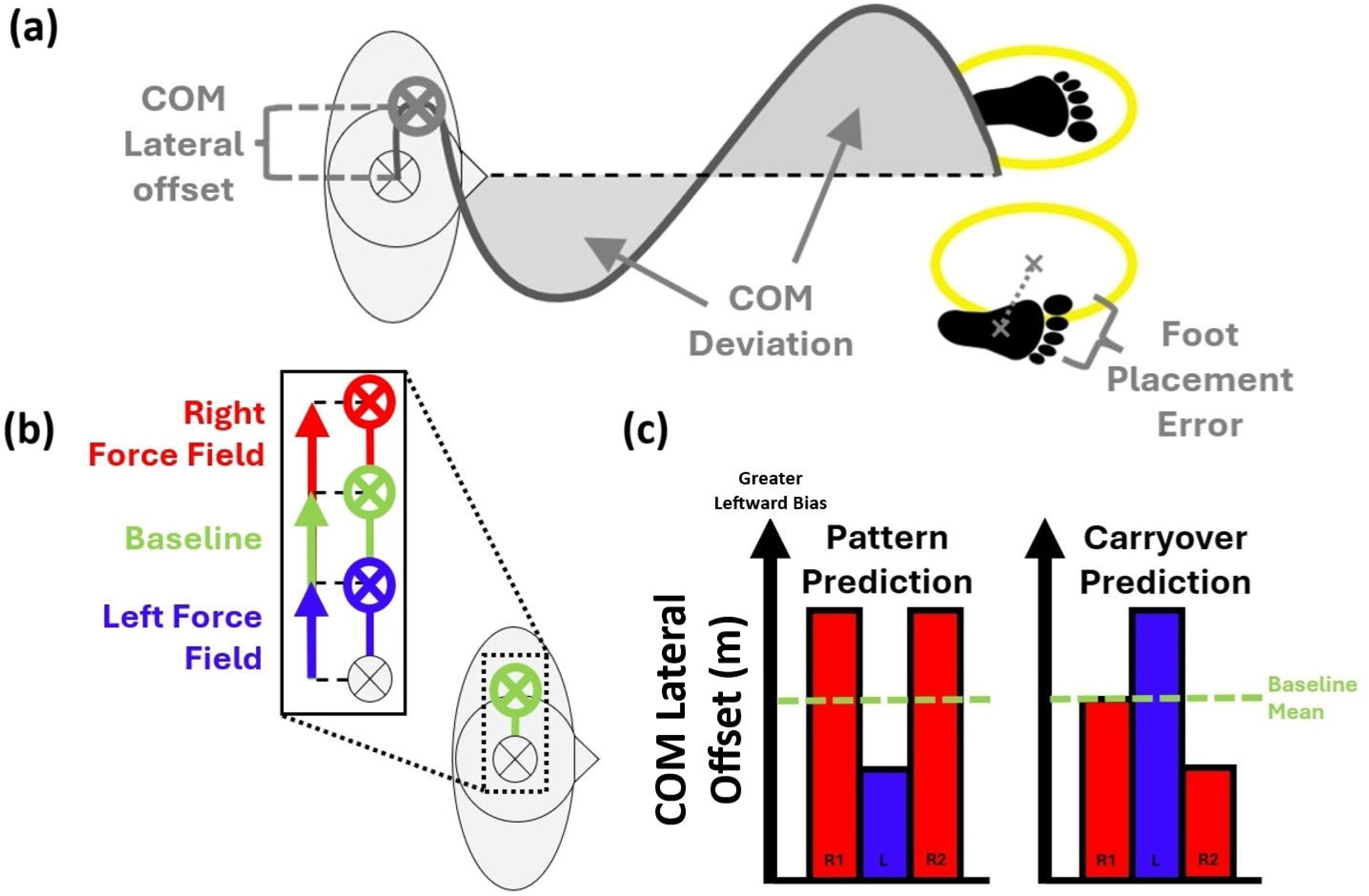
(a) Performance metrics. See **2.5 Data Processing** for more detail. (b) Anticipatory modulation of COM lateral offset. From prior research (Bucklin et al., 2019, 2023; Jiang & Grover et al., 2024), participants increase leftward bias to predictively counter Right Force Fields and decrease it to predictively counter Left Force Fields. (c) Predictive strategy hypotheses operationalized for COM lateral offset during the Triplet Trials. Pattern prediction: high COM lateral offset on R1 and R2, low on L, reflecting accurate prediction of full pattern. Carryover prediction: high on L, low on R2, reflecting (inaccurate) predictions that L is a Right trial, and R2 is a left trial.

Table 1 summarizes the hypotheses for each group. We expected the **Consistent-Certain** group to show **pattern prediction**, reflecting accurate anticipation of Force Field direction even on Switches. We likewise expected the **Inconsistent-Certain** group to exhibit **pattern prediction**, though possibly to a lesser degree as frequent Switches could interfere with learning the specific strategy for each direction. Following Bucklin et al. (2023), we expected the **Consistent-Uncertain** group to exhibit **carryover prediction**, owing to the infrequent but uncertain Switches. In the **Inconsistent-Uncertain** group, however, we expected that the high degree of uncertain Switches would lead participants to avoid predictive strategies altogether, possibly due to pursuing less cognitively demanding reactive or protective strategies.

**Table 1.** Hypotheses – Expected Strategies and Adaptation Rates.

| Group | Predictive Strategy? | Expected Adaptation Rate |
| --- | --- | --- |
| Consistent-Certain | Pattern Prediction | Greatest |
| Inconsistent-Certain | Pattern Prediction | No specific prediction |
| Consistent-Uncertain | Carryover Prediction | No specific prediction |
| Inconsistent-Uncertain | None | Worst |

We further hypothesized that, due to reliance on predictive strategies, **Consistent** environments would yield **higher sensitivity to Streaks and Switches** (error reductions on Streaks; error increases on Switches), while **Inconsistent environments** would show more gradual error reduction with **less sensitivity to Streaks and Switches**.

Finally, we expected that stronger predictive strategies would yield greater overall error reduction (Jiang & Grover et al., 2024). Thus, the **Consistent-Certain group** was expected to show the **largest error reductions**, and the **Inconsistent-Uncertain group** to show the **smallest error reductions**. Aside from these boundary conditions, we had no hypotheses concerning the relative performance rates of the other two groups.

## 2. Methods

### 2.1 Participants

Forty adults (25 females, 15 males, 28.69 ± 6.20 years of age, 67.24 ± 11.45 kg mass, 164.85 ± 28.14 cm height; mean ± SD) provided informed written consent and participated in the study. All protocols were approved by the Northwestern University Institutional Review Board. Participants self-reported that they were free of any musculoskeletal, neural, and/or vestibular pathologies affecting gait or balance and were able to walk continuously for 30 minutes without undue fatigue or health risk. Participants had no prior experience walking in the Force Fields.

### 2.2 Experimental Setup

Participants performed a series of discrete goal-directed stepping trials. Participants began each trial standing with each foot positioned in start targets (see Fig. 1; green ellipses) projected on the walking surface (Hitachi America, Ltd.). Following a start cue (an auditory “beep”), the participant stepped quickly from the start targets to end targets (an identical set of ellipses) located 1.5 × leg length ahead of them. A horizontal line was projected at one third the distance to the end targets, which participants were instructed to always cross with their first step. For safety, participants wore a trunk harness attached to an overhead support device (Aretech LLC, Ashburn VA) that moved freely on a passive trolley in the AP direction. The shoulder straps of the trunk harness were adjusted so that they did not provide bodyweight support or restrict lateral movements.

#### 2.2a Creation of External Walking Environments

During all trials, a cable-driven robotic device applied laterally directed forces to a snug pelvic harness worn by the participant (Fig. 1). Full technical specifications of the robotic apparatus, its pelvic harness attachment, and the application of lateral forces are described in Brown et al. (2017).

Depending on the trial condition, the Agility Trainer produced one of two distinct types of forces: Null Forces or Force Field.

*Null Forces*: The tension in the left and right cables were equal in magnitude resulting in no net force applied to the participant.

*Force Field*: A continuous lateral Force Field directed toward the participant’s right or left side (Fig. 1) proportional in magnitude to their forward walking velocity. Further detail about the Force Field control paradigm can be found in Jiang and Grover et al. (2024), and the same applied Force Field gain (80 N/(m/s)) used in that study and others (Bucklin et al., 2018; 2023) was used for all participants. Maximum applied force across all trials was 18.7 ± 3.2% of participant body weight.

To prevent participants from attempting to “probe” the Force Field direction by slightly leaning forward prior to first toe-off, the Force Field did not onset until participants’ forward movement exceeded a velocity threshold (0.4 m/s). This threshold was identified during previous research (Jiang & Grover et al., 2024) as a velocity during which participants were in single-limb support.

We provided visual feedback to help participants maintain a target peak forward velocity of 1.2 m/s (and thus a consistent Force Field magnitude). A monitor at the end of the walkway displayed “too slow” (<1.1 m/s), “too fast” (>1.3 m/s), or “good job” (within range) (Bucklin et al., 2023; Jiang & Grover et al., 2024).

#### 2.2b Kinematic Measurements

A 12-camera passive motion capture system (Qualisys, Gothenburg Sweden) recording at 100 Hz was used to capture 3D coordinates of reflective markers placed on the participant’s pelvis and feet. Specifically, the pelvis location was tracked using three markers arranged in a triangle cluster over the L5-S1 vertebrae and markers placed bilaterally on the greater trochanter. Foot position was tracked with markers placed on the lateral malleolus, calcaneus, and 2^nd^, 3^rd^, and 5^th^ metatarsals, resulting in a total of 15 markers used for all kinematic measurements.

### 2.3 Protocol

Participants were randomized into four groups of ten: **Consistent-Certain**, **Inconsistent-Certain**, **Consistent-Uncertain**, and **Inconsistent-Uncertain**. Demographic characteristics were similar between groups (see Table 2).

**Table 2.**
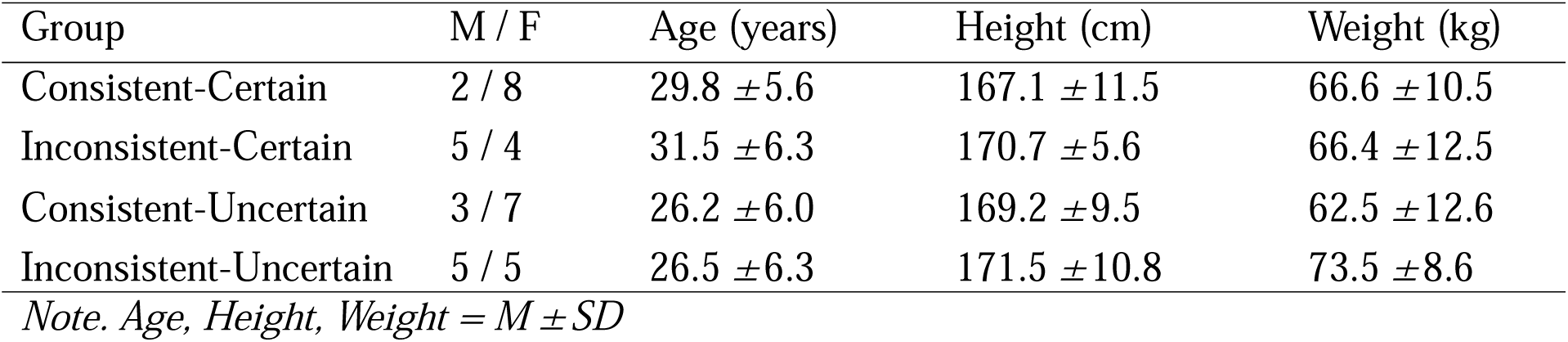
Participant Characteristics.

All participants experienced the same series of trial blocks consisting of 20 Baseline trials (Null Forces), 165 Training Period trials (Force Field), and 20 Washout trials (Null Forces), resulting in 205 total trials (Fig. 1b).

For the Training Period trials, the Force Field’s direction (Right or Left) followed a different schedule per group (Fig 1c). Participants in the **Certain** groups experienced fixed trial schedules; the **Consistent-Certain** group experienced fixed Streaks of five trials in each direction, while the **Inconsistent-Certain** group experienced a fixed Switch sequence of Right and Left trials. Participants in the **Uncertain** groups experienced trials on a probabilistic basis; for the **Consistent-Uncertain** group, each trial had a 90% chance of being a Streak (same direction as previous trial). In contrast, for the **Inconsistent-Uncertain** group, each Force Field trial had a 90% chance of being a Switch (opposite direction from previous trial).

To investigate how the different training groups affected the development of predictive control strategies, all groups experienced 3-trial fixed sequences of Right-Left-Right directed Force Field trials (Triplet Trials) five times, at regular intervals (every thirty trials) within the Training Period (Fig. 1b). The 165 total Training Period trials thus consisted of 150 regular Force Field trials (75 Right and 75 Left), and the five sets of Triplet Trials.

### 2.4 Procedure

Each trial followed the stepping task and setup described in the Experimental Setup section. At the start cue, participants initiated movement with the right foot and completed three steps to reach the end targets. During Training Period trials, the Force Field activated once forward velocity exceeded 0.4 m/s and remained on until the third step. An end cue (same auditory “beep”) then signaled participants to return to the start position. To minimize arm movement confounds, participants crossed their arms during trials but otherwise performed the task in the manner they felt most comfortable.

Before the experimental blocks, participants completed three practice trials with Null Forces to learn the task and then four practice trials with the Force Field (two in each direction) to get an initial sense of its general magnitude.

### 2.5 Data Processing

Kinematic marker data was processed using Visual3D (C-Motion, Germantown, MD) and a custom MATLAB (MathWorks, Natick, MA) program. Marker data was gap-filled and low-pass filtered (Butterworth, 6 Hz cut-off frequency). COM position was calculated in Visual3D as the center of the pelvis model, determined by the three pelvic and two greater trochanter markers.

Figure 2a depicts our outcome measures. We computed **foot placement error** (**FPE**) as our primary (explicit) performance metric. Error for each foot was calculated as the Euclidean distance between the centroid of the foot marker polygon (i.e., the polygon formed by the markers on the calcaneus, 5^th^ metatarsal, and 2^nd^ metatarsal) and the center of the respective end target. Then, overall error was calculated as the root mean square of the errors of both feet.

We characterized control of COM position as a secondary (implicit) performance metric. We analyzed kinematic data of the COM trajectory between the start and end targets and calculated the overall lateral **COM deviation** during each trial (Fig. 2a). This was done by first computing the COM *signed* deviation, which is the signed integral of the COM trajectory, assessed between the moments of the start and end cues, relative to a straight-line path originating from the COM position during the start cue. We normalized COM Signed Deviation relative to Baseline by subtracting the averaged COM Signed deviation taken over the last four Baseline trials^1^. Finally, we normalized for direction (after normalizing to baseline) by taking the absolute value. COM deviation was thus an error metric: greater deviations in COM trajectory (via differences between areas on either side of the straight path) led to larger COM deviation values and indicated greater disruption to the walking trajectory when compared to unperturbed walking.

To gain insight into the strategies underlying the control of COM trajectory, we evaluated lateral movement of the COM prior to first toe-off (**COM lateral offset**) as a potential predictive strategy (Fig. 2a) as this movement always preceded the onset of the directional Force Field. We calculated COM lateral offset as the lateral distance between the COM position during quiet standing (the moment of the start cue) and the maximum leftward excursion of the COM position when unloading the right foot to enable toe-off of the first step.

### 2.6 Analyses

We assessed changes in performance metrics (**FPE, COM deviation**, **COM lateral offset**) in three ways. First, we conducted preliminary assessments of basic task performance across four periods: **Baseline** (average of last four Baseline trials), **Early Field** (first four Training trials, any direction), **Late Field** (last four Training trials, any direction), and **Washout** (last four Washout trials). These periods established baseline performance, initial response to the Force Field, adaptation by the end of the Training Period, and recovery back to baseline behavior.

Second, we quantified how participants adapted to Switches and Streaks in Force Field direction across the full set of Training Period trials (see Statistical Analysis below).

Finally, our primary objective was to test for predictive strategies. To investigate this, we quantified changes in COM lateral offset (as a predictive strategy; see Fig. 2b) across each of the Triplet Trial sets. Trials within each triplet are referred to as **R1** (first Right), **L** (Left), and **R2** (second Right).

Figure 2b depicts how changes in COM lateral offset relative to baseline reflect an anticipation of Force Field direction, as found in previous studies (e.g., Bucklin et al., 2019; Bucklin et al., 2023; Jiang & Grover et al., 2024). Increased offset (greater leftward bias) preemptively resists Right Force Fields, while decreased offset resists Left Force Fields.

Figure 2c depicts how the hypotheses from Table 1 were operationalized for COM lateral offset. Changes that correctly anticipate each change in Force Field direction would indicate pattern prediction. Pattern prediction would therefore be indicated by high COM lateral offset values for R1 and R2, and low values for L Carryover prediction would be reflected in high COM lateral offset on L and low offset on R2, consistent with expecting each trial to match the previous one. R1 would average near baseline, since the preceding (non-triplet) trial was equally likely to be Right or Left.

### 2.7 Statistical Analysis

#### 2.7a Preliminary Assessments

For the preliminary assessments of task performance metrics (FPE and COM deviation), each were submitted to a one-way repeated-measures ANOVA to evaluate overall change across the four experimental periods: Baseline, Early Field, Late Field, and Washout.

#### 2.7b Hierarchical Linear Modeling — Adaptation to Force Field Trials

To assess how participants adapted to the changing Force Field directions over the Training Period, we used hierarchical linear modeling (HLM). This analysis was chosen to preserve the structure of individual adaptation curves and employed an autoregressive variance-covariance structure in the HLM to account for the temporal dependence among trials in the Training Period (i.e., history effects of training).

To consider how Switches and Streaks in Force Field direction affected error over training, we constructed a dummy-coded variable, **Switch-or-Streak**. Each training trial was coded as 1 if it was a Switch (previous trial was opposite), and as 0 if it was a Streak (previous trial was the same).

The **Switch-or-Streak** variable was imbalanced, as each Consistency group had uneven frequencies of Switch and Streak trials (the Consistent groups predominantly had Streak trials, and the Inconsistent groups predominantly had Switch trials). To account for this imbalance, we implemented a participant-level nonparametric bootstrap to estimate uncertainty of fixed-effect coefficients. In each of 1,000 bootstrap iterations, participants were resampled with replacement, retaining all associated trial data. Each resampled participant was assigned a unique identifier to preserve the random intercept structure, the HLM was re-fit, and fixed-effect coefficients were stored. From the resulting empirical sampling distributions, we estimated 95% confidence intervals (CIs) for each coefficient. Coefficients whose intervals excluded zero were considered significant at *α* = 0.05. This bootstrap approach enabled inference while avoiding Type I errors that could result from treating trials as independent or relying only on model-based standard errors under data imbalance.

The final model tested fixed effects of **Switch-or-Streak**, **Trial Number**, **Consistency**, and **Certainty**, plus their interactions. We used a backwards stepwise approach, removing non-significant higher-order terms only when doing so did not reduce model fit (tested by change in -2 log likelihood, chi-square distributed with degrees of freedom equal to the difference in parameters between nested models). To capture nonlinear adaptation curves, we also considered quadratic and logarithmic transformations of Trial Number.

#### 2.7c Hierarchical Linear Modeling — Predictive Strategies

For our primary objective, HLM was used to evaluate if participants exhibited directional bias in COM lateral offset during the Triplet Trials. We focused our analysis on 1) how COM lateral offset changed within a triplet, and 2) if this change within triplets was affected by training. To examine the effects of training, we looked at the first triplet (Early Training) and the last triplet (Late Training). This resulted in a 2 (R1-L or R1-R2) × 2 (Early vs. Late Training) design and was performed separately for each individual group and for each triplet comparison (R1-L and R1-R2).

For all statistics, Greenhouse-Geisser corrections were applied wherever the assumption of sphericity was violated. When the assumption of equal variances was violated for between-subjects comparisons, unpaired comparisons with equal variances not assumed were performed instead. Holm-corrected post-hoc tests and simple effects analyses were run to follow up on significant effects. All statistical tests used *α* = 0.05.

## Results

### 3.1 Preliminary Assessments

All participants complied with the experimental instructions and were able to perform the experiment without falls, reported fatigue, or reported discomfort.

Broadly, participants in all groups experienced significant disruptions to their balance at the start of the Training Period, affecting both explicit (FPE) and implicit (COM deviation) task performance, but then adapted to the Force Field Trials over time (see Fig. 3). Mixed ANOVA indicated that FPE and COM deviation changed as a function of experimental period (*F*(2.14, 76.95) = 84.67, *p* < 0.001, *η_p_^2^* = 0.70, and *F*(2.26, 81.33) = 64.73, *p* < .001, *η_p_^2^* = 0.64, respectively). FPE increased from Baseline (0.02 ± 0.02 m) to Early Field (0.15 ± 0.06 m), *d* = 2.97, *p* < .001, and COM deviations increased from Baseline (0.01 ± 0.004 m^2^) to Early Field (0.08 ± 0.04 m^2^), *d* = 2.67, *p* < .001, as well, indicating an initial disruption to balance and task performance.

**Figure 3.**
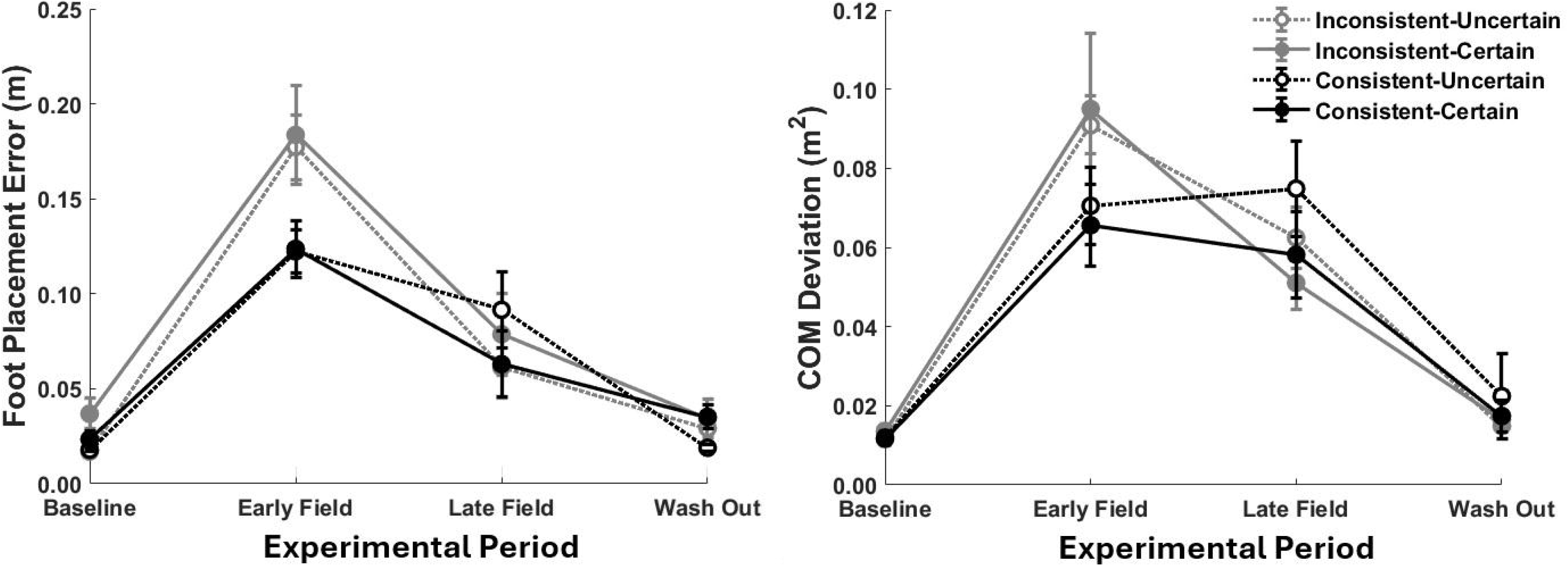
Basic Task Performance—averages of FPE (left) and COM deviation (right) across the four experimental periods.

Participants adapted to the Force Field trials across all groups. FPE decreased from Early Field to Late Field (0.07 ± 0.06 m), *d* = 1.93, *p* < .001, and COM deviation decreased from Early Field to Late Field (0.05 ± 0.03 m^2^), *d* = 1.12, *p* < .001. However, in Late Field, FPE and COM deviation remained greater than Baseline (*d* =1.04, *p* < .001, and *d* = 1.55, *p* < .001, respectively), indicating that participants did not develop Baseline levels of performance during the Force Field trials.

There were decreases in FPE across all groups from Late Field to Washout (0.03 m ± 0.02 m), *d* = 0.91, *p* < .001, and COM deviation from Late Field to Washout (0.0 ± 0.04 m^2^), *d* = 2.67, *p* < .001. There were no differences between Baseline and Washout for either measure, all *p* > .05, indicating participants had returned to Baseline stepping behaviors.

Interestingly, for FPE, there was a two-way interaction between Consistency and experimental period, *F*(2.32,83.51)=3.48, *p* < .05, η²ₚ = 0.09. Simple effects analysis indicated that participants in the Inconsistent Groups exhibited greater FPE in the Early Field (0.18 +/- 0.07 m) than the Consistent Groups (0.12 +/- 0.04 m), *F*(1,38) = 10.57, *p* < .01. There were no other differences between groups at any experimental period, for FPE or COM deviation, all *p* > .05.

#### 3.2a Adaptation to Force Field Trials — Foot Placement Error

FPE data from representative individuals are shown in Figure 4. Visual inspection of these data suggested that, under Consistent conditions, regardless of Certainty, participants were highly sensitive to Switches or Streaks in Force Field direction. Streaks in Force Field direction appeared to precipitate steep declines in FPE, and Switches in direction appeared to precipitate large jumps in FPE, both early and late in training.

**Figure 4.**
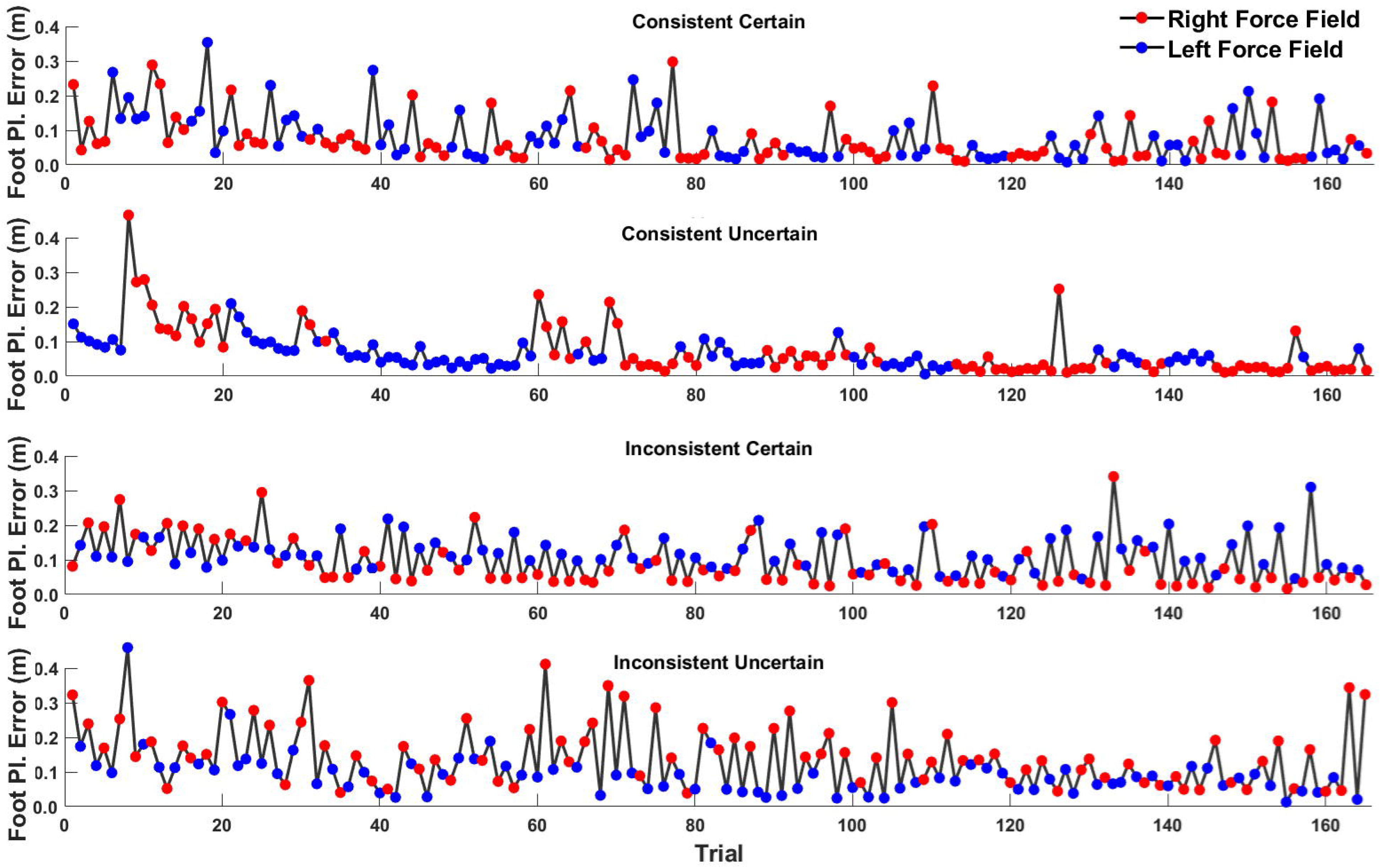
FPE data from one representative subject in each experimental group. Regardless of Certainty, FPE appeared to decrease predominantly during Streaks and remain high during Switches, though a gradual decrease could be seen in the Inconsistent conditions.

The HLM revealed a logarithmic relationship between Trial and FPE. The full model, testing all four predictors—log(Trial), Switch-or-Streak, Consistency, and Certainty—indicated a significant four-way interaction, *β* = -0.03 m, 95% CI [-0.06 -0.01]. There were no other significant effects aside from a fixed main effect of Consistency, *β* = -0.066 m, 95% CI [-0.15 -2e-4], and fixed effect of log(Trial), *β* = -0.023 m, 95% CI [-0.041 -0.012].

Visual inspection of the representative data from individual subjects (such as shown in Fig. 4) suggested the four-way interaction may be driven chiefly by Consistency, where general error reduction was modified by jumps in error during Switches in trial direction, but only for the Consistent groups. As such, we split the model by Consistency and ran HLM separately for the Consistent and Inconsistent groups. For the Consistent groups, HLM indicated a three-way interaction between Certainty, log(Trial), and Switch-or-Streak, *β* = -0.021 m, 95% CI [-0.039 -0.003].

Splitting this data by Certainty, HLM indicated for the Consistent-Certain group an interaction between Switch-or-Streak and log(Trial), *β* = -0.02 m, 95% CI [-0.04 -0.02]. In the model, log(Trial)’s main effect coefficient carried a negative sign, which indicates general error reduction, and the Switch-or-Streak’s main effect coefficient carried a positive sign, which indicates FPE increased on Switches (coded as 1) and decreased during Streaks (coded as 0). Thus, the negative sign of their interaction term indicates that the effect of Switch-or-Streak on FPE diminished as training went on. Conversely, it also indicates that general error reduction was disrupted by Switches and facilitated by Streaks. For the Consistent-Uncertain group, HLM likewise indicated fixed effects of log(Trial), *β* = -0.01 m, 95% CI [-0.01 -0.003], indicating general learning, and Switch-or-Streak, *β* = 0.08 m, 95% CI [0.01 0.15], indicating jumps in error on Switches. However, there was no interaction between Switch-or-Streak and log(Trial) (95% CI crossing zero), indicating that the effect of Switch-or-Streak did not change with learning.

For the Inconsistent groups, HLM indicated no three-way interaction between Certainty, log(Trial), and Switch-or-Streak, nor any effect of Certainty or Switch-or-Streak at any level (all 95% CI crossing zero). There was only a fixed effect of log(Trial), *β* = -0.025 m, 95% CI [-0.043 -0.012], indicating general learning with training.

#### 3.2b Adaptation to Force Field Trials — Center of Mass Deviation

To provide a basic impression of error adaptation over training, COM deviation data from a representative individual are shown in Figure 5. Visual inspection of these data suggested similar trends as for FPE—under Consistent conditions, regardless of Certainty, participant error was highly sensitive to switching or streaking in Force Field direction. Streaks in Force Field direction appeared to precipitate steep declines in COM deviation, and Switches in direction appeared to precipitate large jumps in COM deviation, both early and later in training.

**Figure 5.**
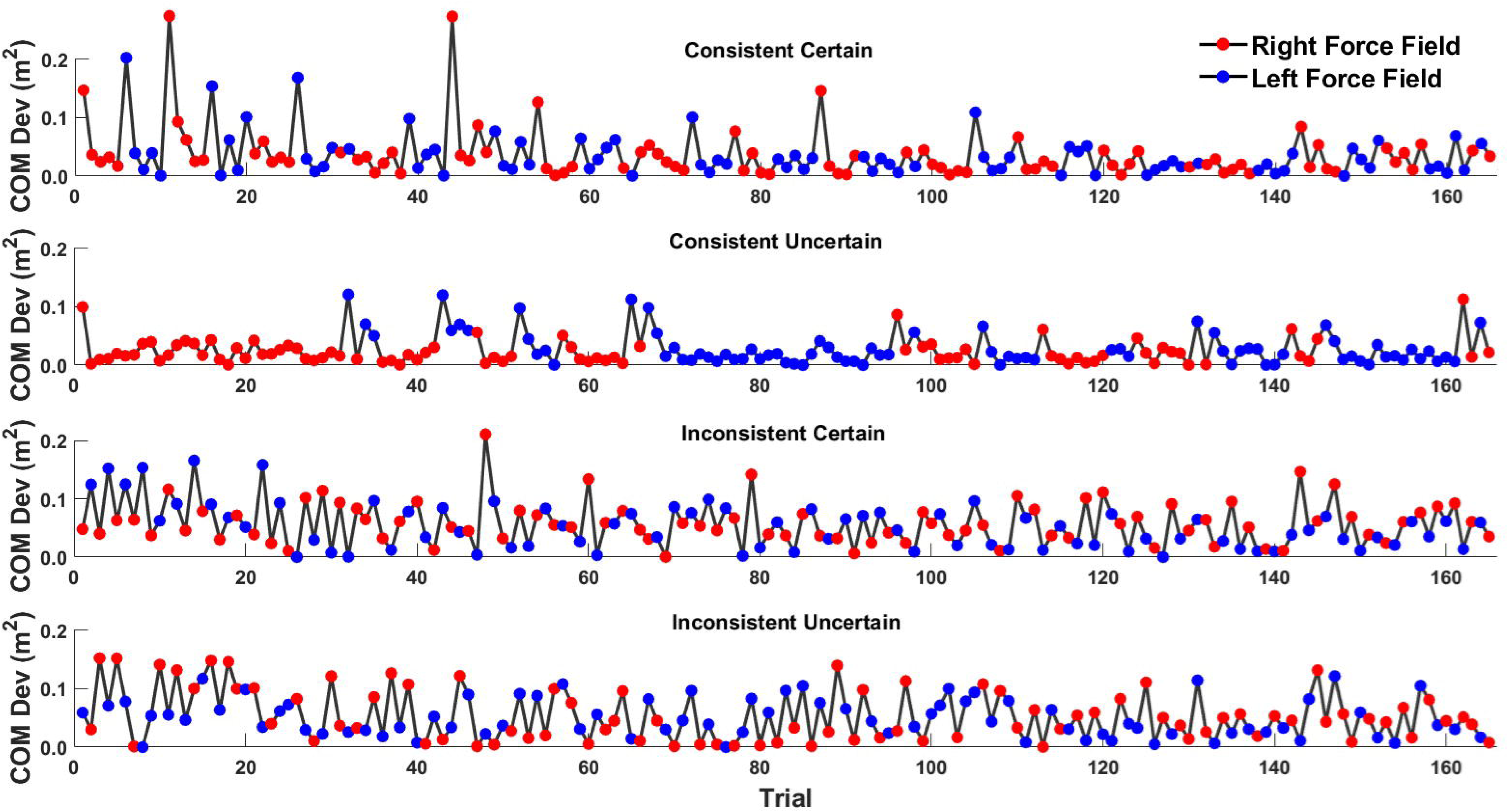
COM deviation data from one representative subject in each experimental group. COM deviation appeared to followed similar trends as FPE.

COM deviation also showed a logarithmic relationship with Trial and there were no four-way interactions, 95% CI crossing zero. There was, however, a three-way interaction between log(Trial), Switch-or-Streak, and Consistency, *β* = -0.009 m^2^, 95% CI [-0.016 -0.0019]. Following our same route for assessing FPE, we split the model by Consistency and ran HLM separately for the Consistent and Inconsistent groups.

For the Consistent groups, HLM indicated a significant two-way interaction between log(Trial) and Switch-or-Streak, *β* = -0.011 m^2^, 95% CI [-0.016 -0.006]. As with FPE, the negative sign of this interaction indicates that the positive effect of Switch-or-Streak (COM deviation increasing on Switches and decreasing during Streaks) diminished with training. Conversely, it also indicates that general learning (reduction of COM deviation during training) was disrupted by Switches and facilitated by Streaks.

For the Inconsistent groups, HLM revealed only a main effect of log(Trial), *β* = -0.011 m^2^, 95% CI [-0.016 -0.006], indicating general learning with training.

For both Consistent and Inconsistent groups, there was no effect of Certainty at any level on COM deviation during training (all 95% CI crossing zero).

### 3.3 Predictive Strategies – COM lateral offset

We investigated whether participants developed predictive strategies with training as indicated by a change in COM lateral offset during Triplet Trials (i.e., scheduled Switches in Force Field direction). HLM was performed for each individual Certainty × Consistency group, and for each Triplet Trial comparison in each group (R1-L and R1-R2). See Table 1 in the Introduction for the analyses’ hypotheses.

COM lateral offset results are depicted in Figure 6. The horizontal dashed line in Figure 6 indicates the Baseline average across all participants as a reference.

**Figure 6.**
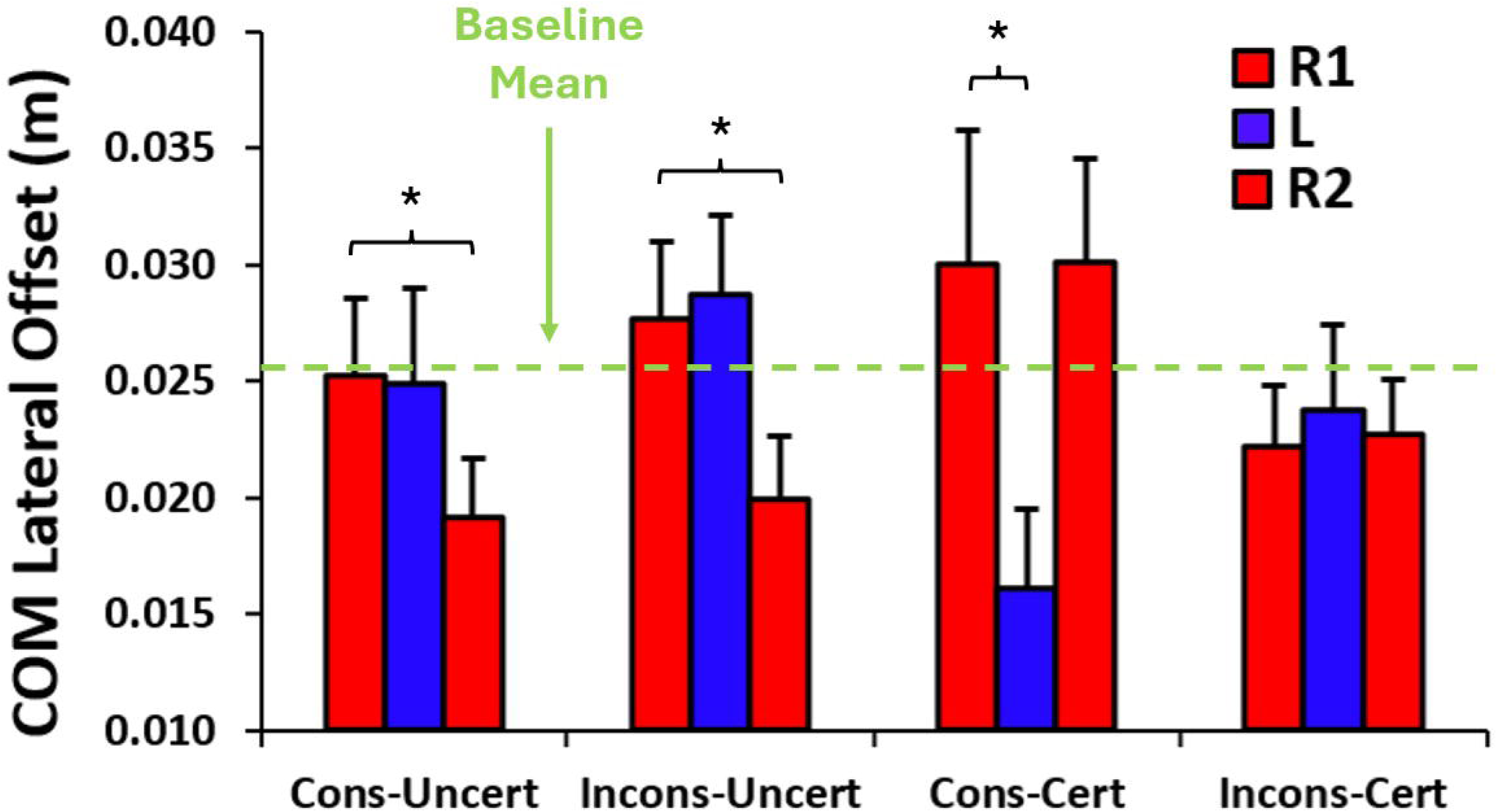
COM lateral offset results (group means ± SE across the first and last sets of Triplet Trials). As expected, the Consistent-Certain group showed COM lateral offset patterns consistent with our hypothesis for pattern prediction (high R1/R2, low L), and the Consistent-Uncertain group showed patterns consistent with our hypothesis for carryover prediction (drop from R1 to R2; see Fig. 2c). Surprisingly, the Inconsistent-Uncertain group appeared to exhibit carryover prediction as well, and the Inconsistent-Certain group exhibited no evidence of predictive strategies. * p < .05

For the Consistent-Uncertain group, HLM indicated for R1-L no differences between Triplet Trial or Training, all *p* > .05. However, there was a main effect of Triplet Trial for R1-R2, *t*(37) = 2.18, *p* < .05. COM lateral offset was lower for R2 (0.019 ± 0.012 m) than for R1 (0.025 ± 0.015 m). For R1-R2 there was no effect of Training nor an interaction between Training and Triplet Trial, both *p* > .05.

For the Inconsistent-Uncertain group, HLM indicated for R1-L no differences between Triplet Trial or Training, all *p* > .05. For R1-R2, there was a main effect of Triplet Trial, *t*(37) = 2.39, *p* < .05. COM lateral offset was lower for R2 (0.020 ± 0.012 m) than for R1 (0.028 ± 0.015 m). For R1-R2 there was no effect of Training nor an interaction between Training and Triplet Trial, both *p* > .05.

For the Consistent-Certain group, HLM indicated a main effect of Triplet Trial for R1-L, *t*(36) = 2.90, *p* < .01. COM lateral offset was lower for L (0.016 ± 0.015 m) than for R1 (0.032 ± 0.016 m). For R1-L there was no effect of Training nor an interaction between Training and Triplet Trial, both *p* > .05. For R1-R2, there were no differences between Triplet Trial or Training, all *p* > .05.

For the Inconsistent-Certain group, HLM indicated no differences between Triplet Trial or Training for either R1-L or R1-R2, all *p* > .05.

## 4. Discussion

The current examined how people adapted during goal-directed stepping when Force Field direction (Left or Right) varied by consistency (Streaks vs. Switches) and certainty (fixed and known vs. probabilistic and unknown). We investigated whether people adopted predictive strategies—pattern prediction or carryover prediction—and how these related to overall error reduction.

We hypothesized predictive strategies (and greatest error reduction) would emerge in consistent and/or certain environments—pattern prediction in the Certain groups (regardless of Consistency) and carryover prediction in the Consistent-Uncertain group. In the Inconsistent-Uncertain group, we expected little prediction and poorer adaptation (see Table 1 and Fig. 2c).

The results partially supported our hypotheses. As expected, participants in consistent environments showed strong sensitivity to Switches and Streaks, consistent with predictive control. Pattern prediction emerged in the Consistent-Certain group, and carryover prediction in the Consistent-Uncertain group. Unexpectedly, carryover prediction also appeared in the Inconsistent-Uncertain group, while the Inconsistent-Certain group showed no predictive strategy. In terms of overall adaptation, Consistent groups showed error reduction mainly within Streaks, with little cumulative improvement, whereas Inconsistent groups improved gradually across trials and were less influenced by trial-to-trial changes.

### 4.1 Adaptation to Force Field Trials

The basic performance results validated the task as a sufficient challenge to walking while still permitting adaptation. As in prior work (Bucklin et al., 2019; 2023; Jiang & Grover et al., 2024), all participants showed large initial explicit (FPE) and implicit (COM deviation) motor errors (Fig. 3; Early Field), then reduced errors over training (Late Field), and returned to Baseline levels after the Washout period.

FPE and COM deviation showed that consistent groups showed greater sensitivity to Switches and Streaks. In both Consistent-Certain and Consistent-Uncertain groups, Switches produced marked error increases, while Streaks produced rapid reductions (Fig. 4; top rows). In contrast, Inconsistent groups showed only gradual error decreases over training (Fig. 4; bottom rows). These results supported our hypothesis that consistent environments heighten sensitivity to Switches and Streaks, likely via predictive strategies.

Contrary to our hypothesis, overall learning was not greater in the Consistent and Certain conditions. In Consistent groups, Switch-or-Streak explained much of the variance in FPE, while the main effect of log(Trial) was small. In Inconsistent groups, FPE was unaffected by Switch-or-Streak but showed larger log(Trial) effects, indicating gradual general learning. Thus, Consistent groups appeared to adjust quickly within Streaks and reach a performance ceiling despite high errors on Switches, leaving little room for further gains. Prior studies show that adaptation to a consistent Force Field can occur within 4–8 trials (Bucklin et al., 2018; Jiang & Grover et al., 2024), suggesting such rapid adjustments could occur within a few Streaks.

### 4.2 Predictive Strategies

COM lateral offset during Triplet Trials (R1–L–R2) offered mixed support for our hypotheses about adaptive strategies to counter the Force Field. Trends resembled expectations—for instance, the Consistent-Certain group showed trial-wise changes consistent with pattern prediction (compare Fig. 2c with Fig. 6)—but no single Triplet Trial in any group or training stage significantly deviated from baseline. Still, differences between trials within triplets were observed, suggesting that although overall biases did not reliably diverge from baseline, participants modulated initial movements between trials. These relative changes provide indirect evidence of predictive adaptation in some conditions but not others.

Participants in the Consistent-Certain group appeared to pursue pattern prediction. COM lateral offset fluctuated across the triplet—high on R1, lower on L, and higher again on R2—indicating proactive biasing based on the global trial pattern. This supports our hypothesis and is unsurprising, given that participants were explicitly informed of the schedule. In contrast, the Inconsistent-Certain group showed no significant differences across Triplet Trials at either training stage, failing to support our prediction that certainty alone would elicit pattern prediction (see Section 4.3b).

The Consistent-Uncertain group appeared to adopt carryover prediction. A decrease in COM lateral offset on R2, following L, implied that participants expected R2 to also be a Left trial. Thus, they proactively biased posture using the most recent trial as a cue. This aligns with our hypothesis that, without explicit knowledge of trial direction, consistent conditions promote reliance on carryover prediction.

Surprisingly, the Inconsistent-Uncertain group also showed signs of carryover prediction. Like the Consistent-Uncertain group, they showed decreases in COM lateral offset on R2, suggesting they expected it to be a Left trial. This contradicted our hypothesis that inconsistent and uncertain conditions would discourage predictive strategies (see Section 4.3a).

### 4.3 How Do Participants Adapt to External Environments of Differing Consistency and Certainty?

Taken together, the results suggest participants pursued pattern prediction only when conditions were both consistent and certain. Under uncertainty, they tended toward carryover prediction, even when patterns were inconsistent and least suited for it. Strikingly, in certain but inconsistent conditions, participants avoided both predictive strategies despite explicit knowledge of upcoming Force Field direction.

#### 4.3a Why Do People Rely on (Inappropriate) Carryover Prediction Under Uncertainty?

These results replicate Bucklin et al. (2023), showing that under uncertainty participants default to assuming the next trial will resemble the last—despite this being maladaptive in inconsistent environments.

In upper-limb studies, exposure to inconsistent or uncertain Force Fields (e.g., alternating curl fields) often leads to an “averaging” or “impedance” strategy (Scheidt et al., 2001), where stiffness is increased through co-contraction (Darainy & Ostry, 2008), tuning resistance across directions (Trumbower et al., 2009). This yields movements that are robust (if suboptimal) across all contexts.

That this did not occur here or in Bucklin et al. (2023) suggests such protective strategies (Franklin et al., 2003; Darainy & Ostry, 2008; Heald et al., 2018; Scheidt et al., 2001; Wei, Wert, & Kording, 2010) are less suitable for whole-body adaptation. Walking likely demands multifaceted control beyond simple stiffening, prompting reliance on cognitive heuristics and pattern-detection biases.

Heuristic frameworks (Kahneman, 2011; Kahneman & Tversky, 1979) show that under limited cues, people default to shortcuts such as recency or representativeness (Tversky & Kahneman, 1974). In this task, a “last-trial = next-trial” heuristic may have appeared cognitively efficient despite frequent failure, as the perceived cost of devising more accurate predictions outweighed the perceived benefit.

Similarly, research in statistical learning suggests that humans are deeply inclined to extract patterns from experience, even when those patterns are illusory (Fiser & Aslin, 2001; Griffiths & Tenenbaum, 2006; Tenenbaum et al., 2006). In noisy or volatile environments, this bias often leads to *apophenia*—the perception of regularities where none exist. From this perspective, carryover prediction may reflect the motor system’s stubborn insistence on using proximal experience to scaffold control, even when such reliance is maladaptive.

Together, these findings suggest that under uncertainty, participants prioritize simplicity and immediacy over statistical accuracy. Even in unsupportive environments, the drive to anticipate persists, reflecting a general principle: the motor system, like cognition, often trades optimality for tractability under informational constraints.

#### 4.3b Inconsistent Environments Are More Obstructive to Whole-Body Adaptation Than to Single-Limb Adaptation

The lack of any change in COM lateral offset across Triplet Trials in the Inconsistent-Certain group was equally surprising. Although we anticipated that frequent switching in Force Field direction might hinder the learning of distinct motor responses for each direction, we nonetheless expected predictive strategies to emerge with practice. However, no evidence of such strategies was observed—even by the end of the training period.

Why participants pursued predictive strategies in Consistent-Certain but not Inconsistent-Certain conditions remains unclear, as both groups received explicit information about upcoming disruptions. In upper-limb adaptation, people can learn and express distinct direction-specific strategies in parallel—even under trial-to-trial reversals—when paired with sufficient contextual cues (Howard et al., 2013; Hirashima & Nozaki, 2012). Our contrasting result suggests that de novo whole-body learning may be more susceptible to contextual interference than single-limb control

One possible explanation is task complexity: whole-body adaptations may be harder to develop and refine. However, prior work shows participants adapt to a consistent Force Field in ∼5–8 trials (Bucklin et al., 2019), comparable to upper-limb reaching tasks (∼8 trials; see Fig. 4 of Lackner & Dizio, 1994).

Another possibility is risk The greater threat of falls in gait adaptation may discourage predictive adjustments. Participants may instead default to more conservative strategies, prioritizing minimizing risk of falls over seeking efficient adaptations. This is consistent with theories of risk-aware control (Sanger 2014), which posit that humans weigh both the probability and cost of failure when selecting motor strategies (Dunning et al., 2015). In uncertain gait environments, the potential cost of miscalibration (i.e., an improperly adjusted step that destabilizes balance) may outweigh the benefit of predictive control, leading participants to sacrifice efficiency for robustness—even when they possess reliable information about upcoming perturbations.

#### 4.3c If Not Predictive Strategies, Then What?

Like other groups, the Inconsistent-Certain group showed high initial errors, gradual reduction during training, and delayed Washout recovery. This meets the definition of *adaptation* from Martin et al. (1996; see also Bastian, 2008), and we can therefore conclude that the Inconsistent-Certain group was adapting *in some way* to the Training Period. The question therefore remains what this group’s adaptation entailed, if not predictive adjustments in COM lateral offset. Some possibilities can be considered, though our study could not assess them directly. The first is that participants were adopting some form of protective strategy, either by increasing stiffness in the leg or otherwise increasing ground reaction forces on the first step to passively resist the Force Field in either direction. A second possibility is that participants were adopting some form of reactive strategy—beginning each trial with the same early movements, with a “plan to react” upon encountering the Force Field.

For example, a few participants in both Inconsistent-Certain and Inconsistent-Uncertain groups adopted a default forward step regardless of direction. When pulled rightward, they quickly rotated the foot outward, shifting the base of support laterally and engaging extensors and plantarflexors to resist the perturbation. When pulled leftward, they remained passive, letting the perturbation guide the foot into a crossover step that extended the base of support and aided balance.

Such a reactive strategy might appear counterintuitive for participants in the Certain group, given their explicit knowledge of forthcoming disruptions. However, this behavior may reflect an emphasis not on prediction, but on the feasibility of the underlying movements. Even with full knowledge of upcoming perturbations, relying on rapid, spontaneous adjustments may be a more effective approach than the more cognitively demanding alternative of planning pre-emptive, resistance-based movements, especially given the contextual interference of inconsistent environments on learning multiple solutions in parallel

#### 4.3d Do Predictive Strategies Increase Reliance on Environmental Consistency?

Previous studies (Bucklin et al., 2019; 2023; Jiang & Grover et al., 2024; Shadmehr et al., 2010) show that predictive strategies can cause large errors when environmental conditions change unexpectedly. For example, after prolonged exposure to a consistent Force Field, sudden removal often produces large compensatory deviations in COM.

Intuition might attribute these errors to mismatches between predicted and actual input, implying that prediction is only vulnerable under unexpected change. Our findings challenge this view. In the Consistent-Certain group, participants clearly engaged in predictive strategies and showed strong sensitivity to Switches and Streaks, even with full foreknowledge of all changes. Thus, errors can persist despite accurate expectations.

Notably, this sensitivity was most pronounced early in the Force Field block and diminished with practice—an effect not observed in the Consistent-Uncertain group, where the impact of Switches and Streaks did not change. This divergence implies that the elevated errors observed during early Switch trials in the Consistent-Certain group did not stem from inaccurate predictions of Force Field direction, but rather from nascent motor execution (motor commands that were still being refined through practice).

Nonetheless, environmental consistency itself appears to play a central role. Both Inconsistent-Uncertain and Consistent-Uncertain groups showed carryover prediction, but only the Consistent-Uncertain group showed heightened sensitivity to Switches and Streaks. These results suggest that sensitivity to sudden changes is primarily shaped by the broader pattern of environmental consistency: large increases in sensorimotor error arise chiefly when changing conditions interrupt otherwise consistent ones, regardless of whether these changes are accurately anticipated.

### 4.4 Limitations and Future Directions

Our analysis was limited by an inability to make direct, between-group comparisons of Switch-versus-Streak effects without resorting to bootstrapping methods. This constrained our ability to formally test whether the magnitude of Switch or Streak sensitivity differed across conditions. A practical solution for future studies would be to incorporate scheduled three-trial Streaks alongside scheduled three-trial Switches (Streak Triplets and Switch Triplets), enabling cleaner comparisons without reliance on resampling techniques.

Our broader research goal is to identify when humans selectively pursue not only *predictive* strategies (anticipating and compensating for an expected disruption), but *reactive* strategies (detecting and responding to a discrete unexpected disruption), and *protective* strategies (proactively resisting any possible unspecified disruption). However, the present study was only equipped to evaluate predictive strategies. Future work should aim to develop experimental paradigms and analytic approaches capable of directly measuring active behavioral signatures of reactive and protective control. A forthcoming study (Kenney, 2025; Kenney et al., in prep) is a promising initial step in this direction.

## Footnotes

1 It was necessary to first take the signed deviation and then normalize to baseline before taking the absolute value, because the baseline deviation carried a slight rightward bias (given that the three-step maneuver always started and ended with a right foot step).

